# The ecological cocktail party: Measuring brain activity during an auditory oddball task with background noise

**DOI:** 10.1101/371435

**Authors:** Joanna E. M. Scanlon, Danielle L. Cormier, Kimberley A. Townsend, Jonathan W.P. Kuziek, Kyle E. Mathewson

**Author notes:** **Corresponding Author:** Kyle E. Mathewson, Ph.D., Department of Psychology, P-217 Biological Sciences Building, University of Alberta, Edmonton, AB, Canada, T6G 2E9.

## Abstract

Most experiments using EEG recordings take place in highly isolated and restricted environments, limiting their applicability to real-life scenarios. New technologies for mobile EEG are changing this by allowing EEG recording to take place outside of the laboratory. However, before results from experiments performed outside the laboratory can be fully understood, the effects of ecological stimuli on brain activity during cognitive tasks must be examined. In this experiment, participants performed an auditory oddball task while also listening to concurrent background noises of silence, white noise and outdoor ecological sounds, as well as a condition in which the tones themselves were at a low volume. We found a significantly increased N1 and decreased P2 when participants performed the task with outdoor sounds and white noise in the background, with the largest differences in the outdoor sound condition. This modulation in the N1 and P2 replicates what we have previously found outside while people ride bicycles (Scanlon et al., 2017b). No behavioural differences were found in response to the target tones. We interpret these modulations in early ERPs as indicative of sensory filtering of background sounds, and that ecologically valid sounds require more filtering than synthetic sounds. Our results reveal that much of what we understand about the brain will need to be updated as we step outside the lab.

## 1. Introduction

Historically, experiments in cognitive neuroscience tend to require participants to experience highly isolated conditions, for two main reasons. The first is to control any effects that the environment may have on normal brain activity. The second is due to equipment limitations that make recording outside of the laboratory difficult and prone to large degrees of data noise. However since these equipment limitations have often made it difficult or impossible to test brain activity in different environments, the effects of ecologically valid environmental influences on brain activity have received little attention. New technologies are beginning to change this, with new mobile EEG technologies that can be carried almost anywhere (Debener, Minow, Emkes, Gandras, & de Vos, 2012; de Vos, Gandras, & Debener, 2014; Zink, Hunyadi, Van Huffel, and de Vos 2016; Krigolson et al., 2017). At this point, few studies have attempted to determine what effect natural environmental noise may have on brain activity. The present study was conducted to simulate some types of natural sound environments in order to determine the effect of environmental sounds may have on brain activity and the ERP.

Several studies have used mobile EEG to perform experiments in real-life environments. Debener et al. (2012) used a wireless EEG system while participants performed an auditory oddball task while walking outdoors and sitting indoors. They were able to show that P3 amplitude was significantly reduced when generated while walking outdoors compared to while sitting indoors. Zink et al. (2016) had participants performing an auditory oddball task during mobile cycling, stationary pedaling and sitting in a natural outdoor environment. The authors found no difference in P3 amplitude between the stationary cycling and sitting conditions, but demonstrated a near-significantly reduced P3 amplitude while participants were cycling compared to both stationary conditions. Recently in our lab, we had participants both sitting and pedaling on a stationary bicycle inside the lab and found no P3 differences during the auditory oddball task (Scanlon, Sieben, Holyk, & Mathewson, 2017a). Next we had participants performing an auditory oddball task both while cycling outside and while sitting inside. Similar to Zink et al. (2016), we found a significantly reduced P3 during the mobile task of cycling. However we were surprised to find a significantly decreased P2 and increased N1 while participants had been cycling outside (Scanlon, Townsend, Cormier, Kuziek & Mathewson, 2017b). Altogether these studies show that in comparison to non-mobile, quiet laboratory conditions, the P3 in an oddball task tends to be reduced during the mobile tasks like walking or cycling outside, but not during stationary cycling inside, indicating that changes in the P3 were related to changes in the task (e.g. stationary vs. mobile). In contrast, when measured and compared to typical laboratory conditions, the N1 is increased and the P2 is decreased only during outdoor cycling. The possibility that this change came from the participants’ surroundings lead us to the current study, in which we attempt to replicate these effects within the laboratory environment.

As our previous mobile EEG study was primarily exploratory, there are several possible reasons for these changes in the N1 and P2 components of the ERP while participants had been cycling outside (Scanlon, et al., 2017b). In the outdoor condition, participants were also cycling, as opposed to sitting in the indoor condition. This meant that they were experiencing a physical (however sub-aerobic) mobile activity, visual optic flow, as well as auditory stimuli that they did not experience while sitting inside the laboratory. However, the modulations in the N1 and P2 were not observed when participants biked on a stationary bike inside the lab (Scanlon et al., 2017a). Additionally, Scanlon et al. (2017b) found that alpha power was decreased while participants cycled outside, and previous work by Brandt, Jansen & Carbonari (1991) has shown that alpha power may be positively correlated with visual N1 and P2 amplitudes. However, most literature concerning the auditory N1 and P2 components point to these components being mostly affected by extraneous auditory factors, especially for a task in the auditory modality (Näätänen & Picton, 1987).

The N1-P2 complex appears to reflect pre-attentive sound processing in the auditory cortex (Näätänen & Picton, 1987). The P1, N1 and P2 components which make up this complex have been shown to reflect several nearly simultaneous processes originating within or near the primary auditory cortex (Näätänen & Picton, 1987; Wolpaw and Penry, 1975; Wood & Wolpaw, 1982). The auditory N1 is a negative-going waveform with three subcomponents, which peak approximately between 75 and 150 ms after stimulus onset (Luck, 2014). The N1 wave appears to be sensitive to auditory stimulus properties and attention, and is increased when an attended stimulus can be more easily distinguished from other stimuli through physical cues such as pitch or location (Näätänen, 1982, 1992). The P2 component, on the other hand, is a positive-going waveform which occurs approximately 150-275 ms after stimulus onset as the second positive component of the ERP (Dunn, Dunn, Languis & Andrews, 1998). Both the auditory and visual P2 are believed to represent a top-down, higher-order aspect of perceptual processing, which is also modulated by attention (Luck & Hillyard, 1994; Freunberger, Klimesch, Doppelmayr & Höller, 2007; Hackley, Woldorff, & Hillyard, 1990). The auditory P2 in particular was shown to decrease when irrelevant speech sounds were introduced to the background of a speech-listening task (Getzman, Golob & Wascher, 2016). Therefore, it appears that this change in the N1 and P2 components relates to auditory aspects of top down perceptual processing and attention that are altered when participants perform the same task in a rich and noisy environment (Scanlon, et al., 2017b).

Perhaps some aspect of an outdoor environment changes the attentional and sensory processes required to perform an auditory task. This could be due to the overlapping sounds, as one filters unimportant background stimuli to focus on their current task. Conversely, the reason could also be as simple as the possibility that the same headphone task sounded subjectively quieter in an open environment. In the current study, we intend to investigate these possibilities by having participants perform the same headphone auditory oddball task as in previous studies (Scanlon et al., 2017a; Scanlon et al., 2017b) with the following conditions: background outdoor traffic sounds, background white noise, silent background, and silent background with quieter tones. Evidence from the literature leads us to hypothesize that the effect within the N1 and P2 in our previous study was due to a process of filtering out extraneous ambient noise, while focusing in on the relevant task. Therefore we hypothesize that we will be able to most effectively replicate the increase in N1 and decrease in P2 when outdoor sounds are played in the background.

## 2. Methods

### 2.1 Participants

A total of sixteen members of the university community participated in the experiment, however 2 participants were removed due to technical errors and incorrect performance of the task, leaving a total of fourteen participants (Mean age=21.6; Age range=18-26; Sex=5 female). Six participants were members or associated members of The Mathewson Lab at the University of Alberta. All participants were either compensated $10/hr or received course credit for participating in the experiment. Participants all had normal or corrected-to-normal vision with no history of neurological problems. The experimental procedures were approved by the Internal Research Ethics Board of the University of Alberta and participants gave informed consent prior to participating.

### 2.2 Materials

In all conditions, participants were seated in a radio frequency attenuated chamber in front of a 1920 × 1080 pixel ViewPixx/EEG LED monitor running at 120 Hz with simulated-backlight rastering. A fixation cross was presented using a windows 7 PC running Matlab 2012b with the Psychophysics toolbox. Video output was via an Asus Striker GTX760 and audio output was through an Asus Xonar DSX sound card. The participants pressed a button with their right index finger when they heard the high oddball tone. Responses from the button press were marked in the EEG data by the Raspberry Pi 2 model B computer.

### 2.3 Procedure

Each participant completed an auditory oddball task in four conditions: *silent, outside sounds, white noise*, and *silent-low* (Figure 1). For the oddball task, a pair of Sony MDR-E10LP headphones played one of two different frequency tones (either 1500 or 1000 Hz; sampled at 44.1 kHz; two channel; 16-ms duration; 2-ms linear ramp up and down; 65 Db). A pair of Logitech Z130 speakers played the outdoor sounds and white noise clip.

**Figure 1.**
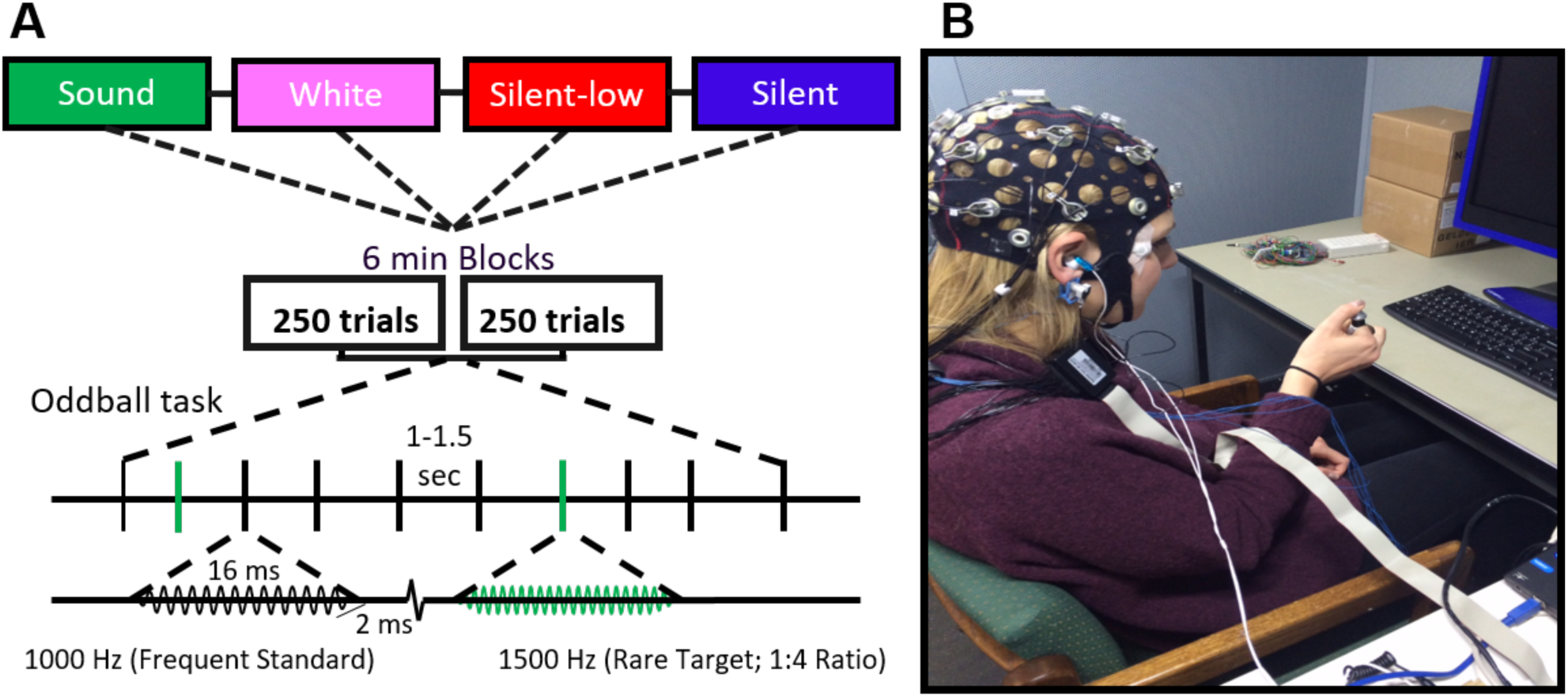
EEG apparatus and procedure. A: During each of the noise conditions, participants performed two 6-minute blocks of an auditory oddball task, with self-paced breaks in between. Within the task, 80% of the tones were frequent low-pitched tones (1000 Hz), and 20% were rare high-pitched tones (1500 Hz). Each tones was played for 16 ms with a ramp-up and down of 2 ms. Tones were played 1-1.5 seconds apart. B: The task was performed in a Faraday cage with the participant wearing an EEG cap and responding to target tones with a hand-held button connected to the Raspberry Pi.

Condition order was counterbalanced across participants and they completed eight 6 minute blocks of 250 trials for a total of 2000 trials. Each trial had a 1/5 likelihood of being a target trial. Each block began with a 10-second countdown to ensure the oddball task only took place when the participant was ready and when the necessary sounds were playing. Each trial also began with a pre-tone interval between 1000 and 1500ms, followed by the tone onset. The next trial began immediately after the tone offset, with participants responding to targets during the following pre-tone interval.

A Raspberry Pi 2 model B computer, which was running version 3.18 of the Raspbian Wheezy operating system, using version 0.24.7 of OpenSesame software (Mathôt, Schreij, & Theeuwes, 2012), was used both to run the oddball task and to mark the EEG for ERP averaging (Kuziek, Shein, and Mathewson, 2017). Audio output was via a 900 MHz quad-core ARM Cortex-A7 CPU connected through a 3.5 mm audio connector. Coincident in time with sound onset, 8-bit TTL pulses were sent to the amplifier by a parallel port connected to the General Purpose Input/Output (GPIO) pins to mark the data for ERP averaging. A start button was affixed to the Raspberry Pi to simplify the beginning of a block. The participant’s task was to press the target button with their right hand when the rare tone was heard while keeping their eyes on a fixation cross.

### 2.4 Conditions

In the *silent* condition the participants were asked to perform the oddball task with no background noise. In the *outside sounds* condition, a 6 minute recording of sounds taken next to a noisy roadway (available upon request) was played in the background, with a maximum volume of 84 dB (Min 55 dB; tones played at 65 dB). The sounds were recorded on an iPhone 5, voice memos app from the same location the previous bike study took place (Scanlon et al., 2017b; bike path along Saskatchewan Dr. NW, Edmonton, Canada; between 116 St and 111 St) in December 2015. In the *white noise* condition an even band of frequencies was played in the background at a volume of 54 dB for 6 minutes. In the *silent-low* condition, no sounds were played in the background, and the oddball task itself was played at a decreased volume of 54 dB.

### 2.5 EEG Recording

Based on Mathewson, Harrison, & Kizuk’s (2015) previous lab work directly comparing overall electrode quality (noise levels, statistical power etc.) active amplified wet electrodes (BrainProducts actiCAP) were selected for the study. Ag/AgCl pin electrodes were used and arranged in 10-20 positions (Fp2, F3, Fz, F4, T7, C3, Cz, C4, T8, P7, P3, Pz, P4, P8, and Oz). Additionally, a ground electrode, embedded in the cap at position Fpz, and two reference electrodes, clipped to the left and right ear, were used (referenced online to the right ear). SuperVisc electrolyte gel and mild abrading of the skin with the blunted syringe tip were used to lower impedances of all the electrodes. Gel application and aforementioned techniques continued until impedances were lowered to < 10 kΩ, measured using an impedance measurement box (BrainProducts) and until data quality appeared clean and reduced of noise. In addition to the 15 EEG sensors, 2 reference electrodes, and the ground electrode, the vertical and horizontal bipolar electrooculogram was recorded from passive Ag/AgCl easycap disk electrodes affixed above and below the left eye, and 1 cm lateral from the outer canthus of each eye. NuPrep preparation gel was applied to the applicable areas of the face, followed by wiping of the skin using an antibacterial sanitizing wipe, both used to lower the impedance of these EOG electrodes based on visual inspection of the data. These bipolar channels were recorded using the AUX ports of the V-amp amplifier, using a pair of BIP2AUX converters, and a separate ground electrode affixed to the central forehead.

EEG was recorded with a Brain Products V-amp 16-channel amplifier. Data were digitized at 500 Hz with a resolution of 24 bits. Data were filtered online, via BrainVision Recorder settings (BrainVision Solutions, Canada), with a 2^nd^ order Butterworth filter and attenuation points at 0.1 and 30Hz (cutoff slope: 12dB/octave). To remain consistent with a previous study which was performed outside of the laboratory (Scanlon et al., 2017b) a notch filter was also applied at 60 Hz. Data was recorded using a Microsoft Surface Pro 3 running Brain Vision Recorder, and powered along with the Raspberry Pi by an Anker Astro Pro2 20000 mAh Multi-Voltage External Battery. The mentioned technology was connected to the Surface Pro 3 using a Vantec 4-Port USB 3.0 Hub.

Trials took place in a dimly lit sound and radio frequency attenuated chamber, with copper mesh covering the window. The fan and lights were turned on, to allow proper ventilation and visual acuity of the fixation. The monitor runs on DC power from outside the chamber, the keyboard and mouse plugged into USB outside the chamber, and the speakers and amplifier were both powered from outside the chamber. Nothing was plugged into the internal power outlets. Any devices transmitting or receiving radio waves (i.e., cellphones) were removed from the chamber for the duration of the experiment.

### 2.5 EEG analysis

EEG analyses were computed using MATLAB 2012b with EEGLAB (Delorme & Makeig, 2004), as well as custom scripts. After recording, EEG data was re-referenced to the average of the left and right ear lobe electrodes. Timing of the TTL pulse was marked in the EEG data during recording, and used to construct 1200-ms epochs (including the 200-ms pretrial baseline) which were time-locked to the onset of standard and target tones. The average voltage in the first 200-ms baseline period was subtracted from the data for each electrode and trial. To remove artifacts caused by movement, amplifier blocking, and any other non-physiological factors, any trials in either of the conditions with a voltage difference from baseline larger than ±1000 µV on any channel (including eye channels) were removed from further analysis. Following eye correction, a second threshold was applied to remove any trials with a voltage difference from baseline of ±500 µV. Over 98 percent of trials were retained after this procedure in each condition. On average, artifact rejection left approximately equal numbers of trials per participant in all conditions (Table 1), from which we computed the remaining analyses.

**Table 1.** Mean, range and standard deviation of trials remaining after artifact rejection in all conditions.

| Condition | Targets |  |  | Standards |  |  |
| --- | --- | --- | --- | --- | --- | --- |
|  | Mean | Range | Standard Deviation | Mean | Range | Standard Deviation |
| **Sound** | 99.929 | 99-100 | 0.267 | 398 | 386-400 | 3.762 |
| **White noise** | 99.5 | 97-100 | 1.092 | 397.1429 | 382-400 | 5.682 |
| **Silent-low** | 99.714 | 97-100 | 0.825 | 397.643 | 378-400 | 6.0461 |
| **Silent** | 99.571 | 96-100 | 1.158 | 398 | 381-400 | 5.054 |

A regression-based eye-movement correction procedure was used to estimate and remove variance due to artifacts in the EEG due to blinks, as well as vertical and horizontal eye movements (Gratton et al., 1983). This technique identifies blinks with a template-based approach, then computes propagation factors such as regression coefficients, predicting the horizontal and vertical eye channel data from the signals at each electrode. This eye channel data is then subtracted from each channel, weighted by these propagation factors, allowing us to remove most variance in the EEG which can be predicted by eye movements. No further rejection or filtering was done on the data in order to include as many trials as possible for all conditions, as well as to investigate how minor sources of non-eye-related noise contribute to the power to measure ERP components during the outdoor cycling task.

### 2.6 Spectral analyses

Spectral analyses were performed on a random sample of 370 of each participant’s artefact-removed standard trials (Figure 4C). The full length (including the 200 ms baseline and 1000 ms after the stimulus) of each of these trials were then used to calculate a fast Fourier transform (FFT) through a procedure of symmetrically padding the 600 time point series with zeros, making a 1,024-point time series for each epoch, providing .488 Hz frequency bins. Because the data were collected with an online low-pass filter of 30 Hz, only frequencies measuring up to 30 Hz were plotted. We then calculated the spectra for each participant by calculating the average of the 370 spectra for each participant, and then combining these into a grand average spectra.

### 2.7 Single trial and ERP RMS analysis

As an estimate of the noise created on single-trial EEG epochs, we calculated the RMS value of a baseline period for each standard trial (de Vos & Debener, 2014). This baseline period consisted of the 100 time points (200 ms) prior to each tone’s onset, in order to avoid any interference of the evoked ERP activity in the measurement of RMS. RMS is equivalent to the average absolute voltage difference around the baseline, and therefore is a good estimate of EEG data single-trial noise. To estimate a distribution of RMS for each condition in our data, we used a permutation test that selects, without replacement, a different set of 360 epochs for each participant’s standard artifact-corrected trials, on each of 10,000 permutations before running second-order statistics (Laszlo et al., 2014; Mathewson et al., 2017).

In order to test for noise that was not effectively averaged out over trials, a similar test was performed on noise levels within trial-averaged ERPs. To quantify the amount of noise in the participant average ERPs, we again used a permutation test of RMS values in the baseline time-window. Complementary to the single-trial analysis, this computation estimates the amount of phase-locked EEG noise in the data that is not averaged out over trials with respect to onset of the tone. We randomly selected and averaged 360 standard trials without replacement from each participant’s standard artifact-corrected trials. The RMS values obtained were then averaged over EEG channels to make a grand average for all participants. This made 10,000 permutations once each participant’s data were averaged together to compute second-order statistics.

### 2.8 Statistical analysis

In this study, we intend to analyze behavioural responses, ERPs and spectral power between all four conditions. Behavioural analysis consists of a one-way repeat measures ANOVA between average oddball task response times in the four sound conditions. ERP analysis for this study will focus on the N1 (Fz; 100-175 ms), P2 (Fz; 175-275 ms), and P3 (Pz; 300-430 ms) ERP components. To assess both condition differences and differences between standards and targets, a 2 × 4 repeat measures ANOVA was conducted for each component with the first factor being standards vs. targets and the second being the four sound conditions. Where significant group differences were found between conditions, pair-wise Bonferroni corrected t-tests were then used to further investigate how the conditions differed within stimulus.

As the task used a button press for response to target tones, it is possible that effects of muscular preparation and activation may be included in the ERP and oscillation effects for targets. Additionally, there is no reason to predict differences between standard and target tones in prestimulus data noise. Therefore, only standard tones were included in the spectral and RMS analysis. For single-trial and baseline ERP RMS data noise a Wilcoxon Signed rank test was used to measure differences between the conditions. A one-way repeat measures ANOVA was used to test for spectral differences between conditions.

## 3. Results

### 3.1 Behavioural differences

A bar graph of the response rate and reaction time is depicted in Figure 2. To test for differences in response rate and average reaction time to targets in each condition, one-way repeat-measures ANOVA test with Greenhouse-Geisser correction was performed, revealing no significant group differences in the response rate (F(3,13) = 0.908, *p* =0.3566) or mean response time (F(3,13) = 1.202, *p* = 0.2779).

**Figure 2.**
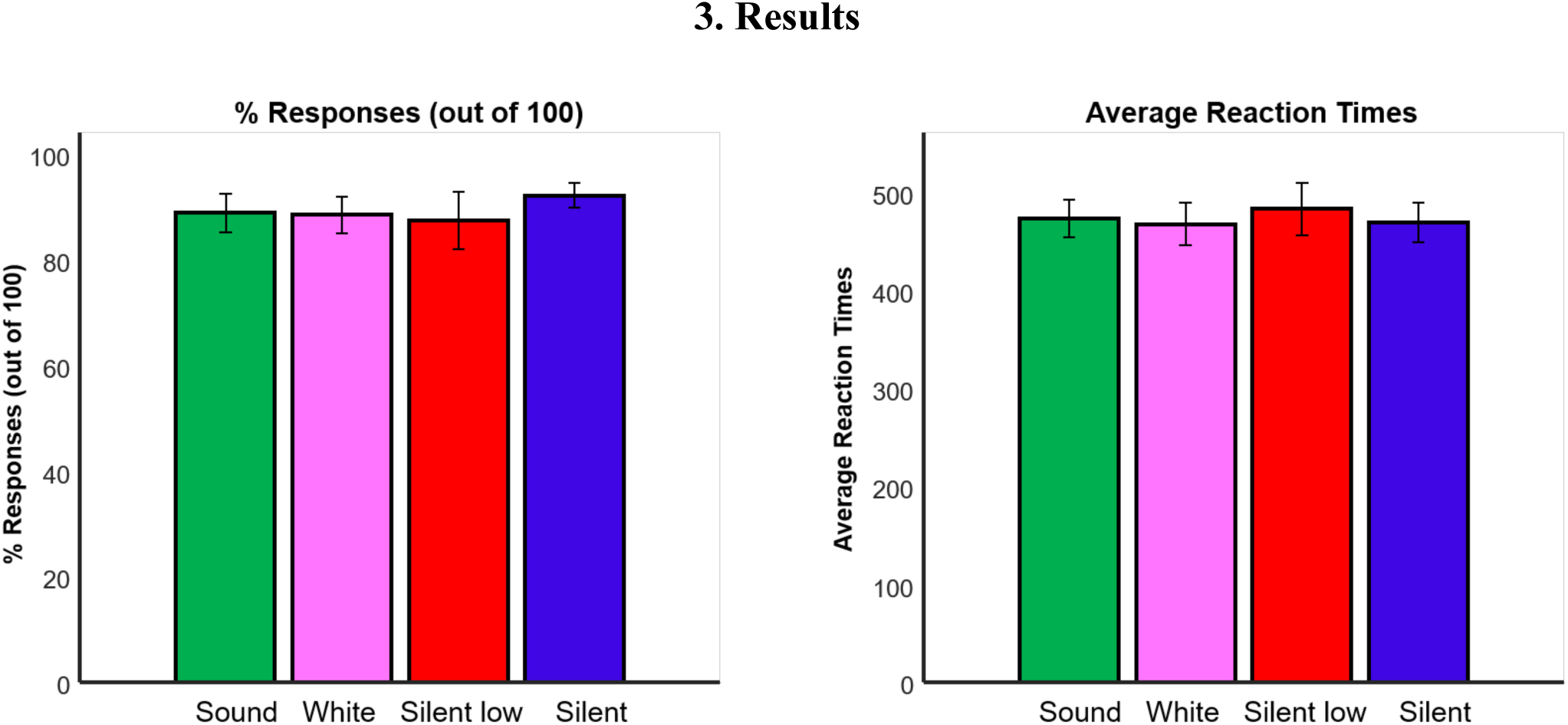
Behavioural analysis. Left: Bar graph depicting mean and standard error for the percentage of targets that received a response before the following tone across participants for both conditions. Right: Bar graph depicting the mean and standard error for the average response time (with missed responses removed) across participants for each condition.

### 3.2 ERP morphology and topography

#### 3.2.1 P3 oddball effects

Figure 3A depicts grand average ERPs calculated from each participant’s artifact-removed and corrected target and standard tones at electrode Pz. Shaded regions represent the standard error of the mean for each time point. Evident from the plots is the expected increase in P3 amplitude during target trials in the central posterior topographical regions (Pz; Figure 3B).

**Figure 3.**
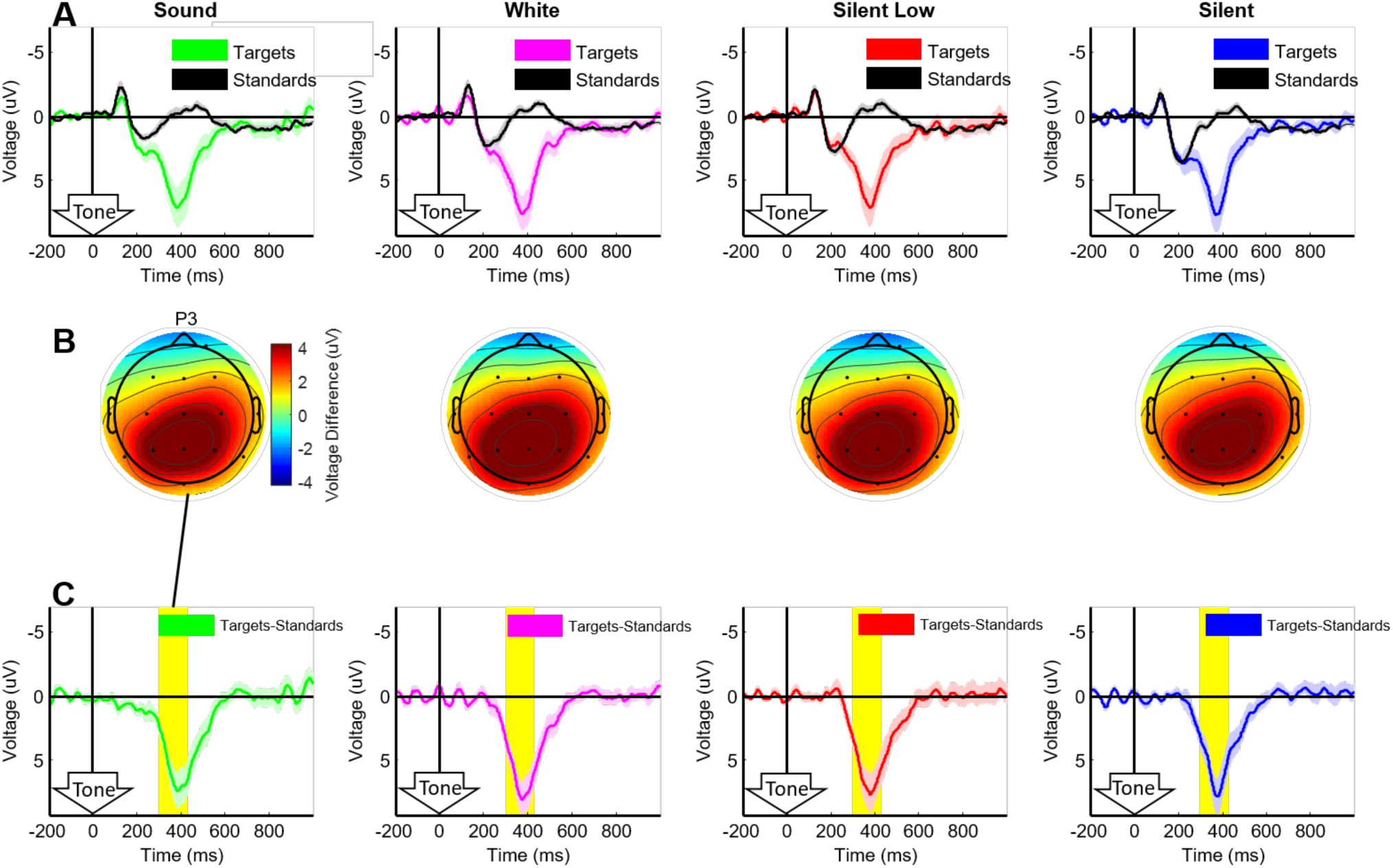
Grand average ERPs. Grand average ERPs. A: Grand-average ERPs computed at electrode Pz for all eye movement and artifact corrected trials, for both target (colour) and standard (black) tones. B: Scalp topographies for grand-average ERP difference between target and standard tones in the P3 time window (highlighted in yellow), 300-430 ms after the tones, respectively. C: ERP difference wave from electrode Pz, with shaded regions representing within-subject standard error of the mean for mean for this difference, with between-subjects differences removed (Loftus & Masson, 1994). Yellow highlighted regions depict the time window for the P3 analysis and topographical plots.

Figure 3B depicts topographies of the target-standard differences within the P3 time window. The P3 topographies demonstrate the expected posterior scalp activation distributions. Figure 3C depicts the ERP difference waves at electrode Pz, which are created through the subtraction of the standard tone ERPs from the target tone ERPs for each subject. Shaded regions depict the within-participant standard error of the mean, with variation between participants removed by a target - standard subtraction. In order to test for significant differences between standard and targets within the P3 time window in all conditions, a 2 × 4 repeat measures ANOVA with Greenhouse-Geisser correction was performed with the first factor being standard and target stimuli, and the second being the four sound conditions. This test revealed a significant effect of stimulus (F(1,13) = 31.675, *p* < 0.001; η_p_^2^ = 0.709) and no effect of condition (F(3,13) = 1.064, *p* = 0.371; η_p_^2^ = 0.076) or interaction (F(3,13) = 1.046, *p* = .508; η_p_^2^= 0.052).

#### 3.2.2 N1 amplitude

In order to observe the effects of the four different sound conditions on general stimulus processing, grand averaged ERPs were separated into standards and targets for each condition at the Fz electrode and plotted in Figure 4B. A visual inspection of the plots in Figure 4B indicates increased amplitude in the N1 component for the *outside sound, white noise* and *silent-low* conditions for the standards and targets at Fz. To test for significant differences for the N1 time window, a 2 × 4 (factors: stimulus and condition) repeated measures ANOVA test with Greenhouse-Geisser correction was used, indicating significant effects of the stimulus (F(1,13) = 13.400, *p* = 0.003; η_p_^2^ = 0.508) and condition (F(3,13) = 8.853, *p* < 0.001; η_p_^2^ = 0.405), but no interaction effects (F(3,13) = 1.059, *p* = 0.377; η_p_^2^ = 0.075).

**Figure 4.**
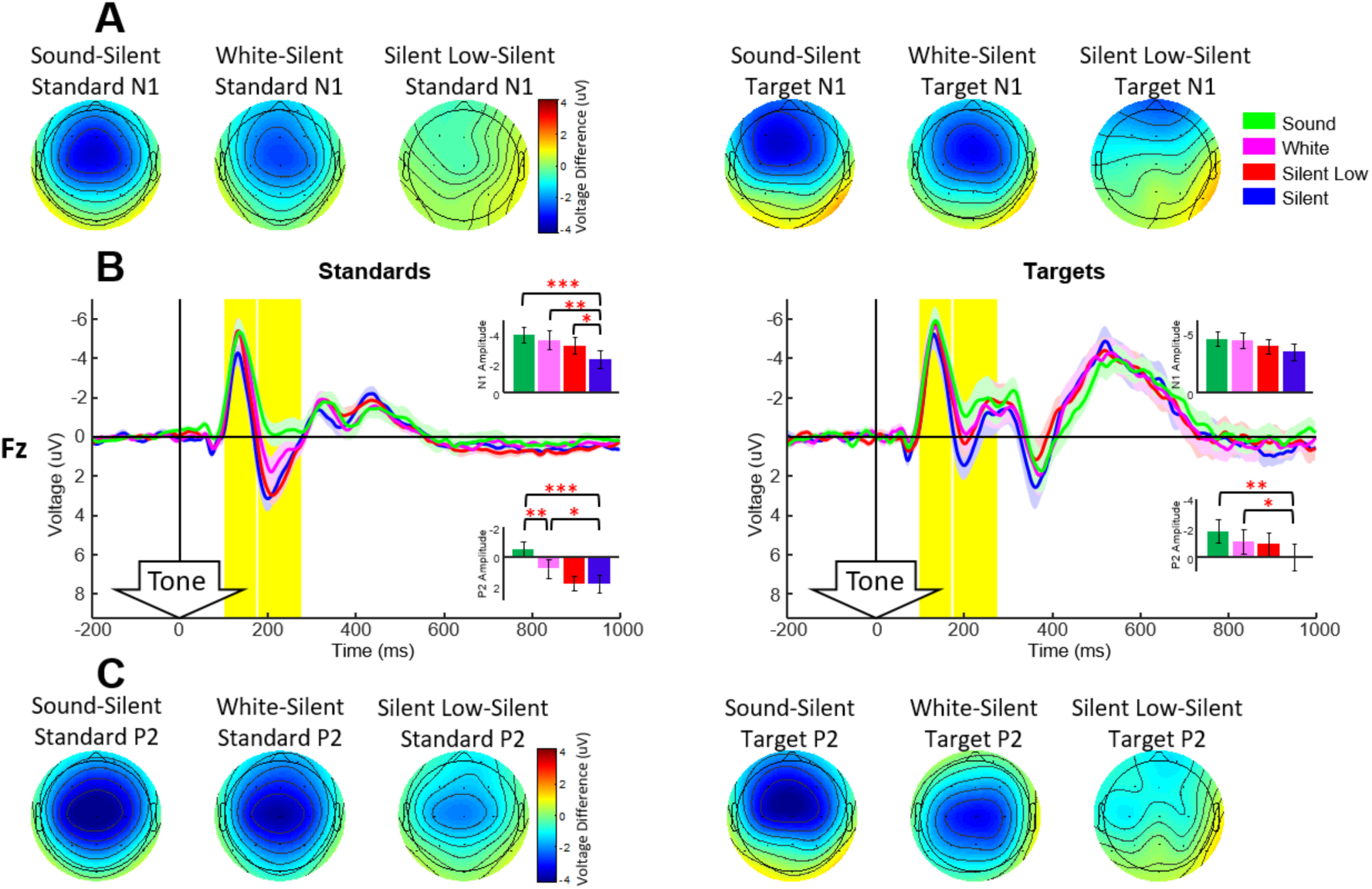
Grand average ERPs A: Topographies for the N1 time window comparing each condition to the silent background condition. B: Grand average ERPs collected at the Fz electrode location, plotted separately to compare within standards and targets between conditions. Shaded regions indicate the standard error of the mean. Inset bar graphs show the mean and standard error across participants in the N1 and P2 time windows. C: Topographies for the P2 time window comparing each condition to the silent background condition. Red asterisks denote significant differences, while green asterisks denote a marginal difference.

In order to further understand the N1 condition effects, four Bonferroni corrected t-tests within both standard and target stimuli (8 altogether; *α* =0.0063)were used to compare each condition to the *silent* condition, as well as comparing the two background-noise conditions. Here we found that the group differences in condition were driven by a difference between the *silent* condition and all three of the sound-altered conditions within standard tones. The *silent* condition was found to have a significantly smaller amplitude N1 compared to the *outside sound* (M_diff_ = -1.683; SD_diff_ = 1.029; *t*(13) = -6.116; *p* = 3.682e-05), *white noise* (M_diff_ = -1.319; SD_diff_= 1.115; *t*(13) = -4.424; *p* = 6.872e-04) and *silent-low* (M_diff_ = -0.901; SD_diff_ = 0.957; *t*(13) = - 3.519; *p* = 0.004) conditions.

#### 3.2.3 P2 amplitude

Visual inspection of Figure 4B at the P2 time window, reveals a higher amplitude positive voltage for the *silent* condition compared to the *sound, white noise* and (for targets) the *silent-low* conditions. To test for significant group differences in the P2 time window at Fz, a 2X4 repeated measures ANOVA (factors: stimulus and condition) with Greenhouse-Geisser correction was performed, revealing significant group differences for stimulus (F(1,13) = 16.521, *p* = 0.001; η_p_^2^ = 0.560), condition (F(3,13) = 21.198, *p* <0.001 ; η_p_^2^ = 0.620) as well as interaction effects (F(3,13) = 3.845, *p* = 0.027; η_p_^2^ = 0.228).

To further investigate the P2 condition effects, four Bonferroni corrected t-tests within both standards and targets (8 altogether; *α* =0.0063)were used to compare each condition to the *silent* condition, as well as comparing the two background-noise conditions. Here we found that these group differences were driven mainly by significant differences between the two background-sound conditions and the *silent* condition in both standard and target stimuli. Within both standard and target stimuli, the P2 amplitude in the *silent* condition had significantly higher amplitude than the *outside sound* (Standards: M_diff_ = -2.348 ; SD_diff_ = 1.2094 ; *t*(13) =-7.2639 ; *p* = 6.327e-06; Targets: M_diff_ = -1.7852 ; SD_diff_=1.558 ; *t*(13) = -4.2872 ; *p* = 8.8e-0402) and *white noise* (Standards: M_diff_=-1.0152 ; SD_diff_=1.1034 ; *t*(13) = -3.4426 ; *p* = 0.0043699; Targets: M_diff_ = -1.0699 ; SD_diff_ = 1.122 ; *t*(13) =-3.5678 ; *p*=0.00344) conditions. Additionally within standard stimuli, the *white noise* condition had significantly higher P2 amplitude than the *outside sound* condition (M_diff_ = -1.3327 ; SD_diff_ =1.0333 ; *t*(13) = -4.826 ; *p*= 3.310e-04).

#### 3.4 Spectral differences

Figure 5C shows the grand average spectra for the Fz and Pz electrodes. Shaded regions indicate the standard error of the mean across participants. Evident from the plots, all four conditions show the expected 1/frequency structure in the data. Also evident are no significant differences in any frequency. To test for differences in alpha power, a one-way repeated measures ANOVA with Greenhouse-Geisser correction was performed in both the Fz and Pz electrode locations, for 9 frequency bins from 7.81 Hz to 11.72 Hz. This test indicated no significant group differences for either the Fz (F(3,13) = 1.351, *p* = 0.254) or Pz (F(3,13) = 0.025, *p* = 0.759) electrodes.

**Figure 5.**
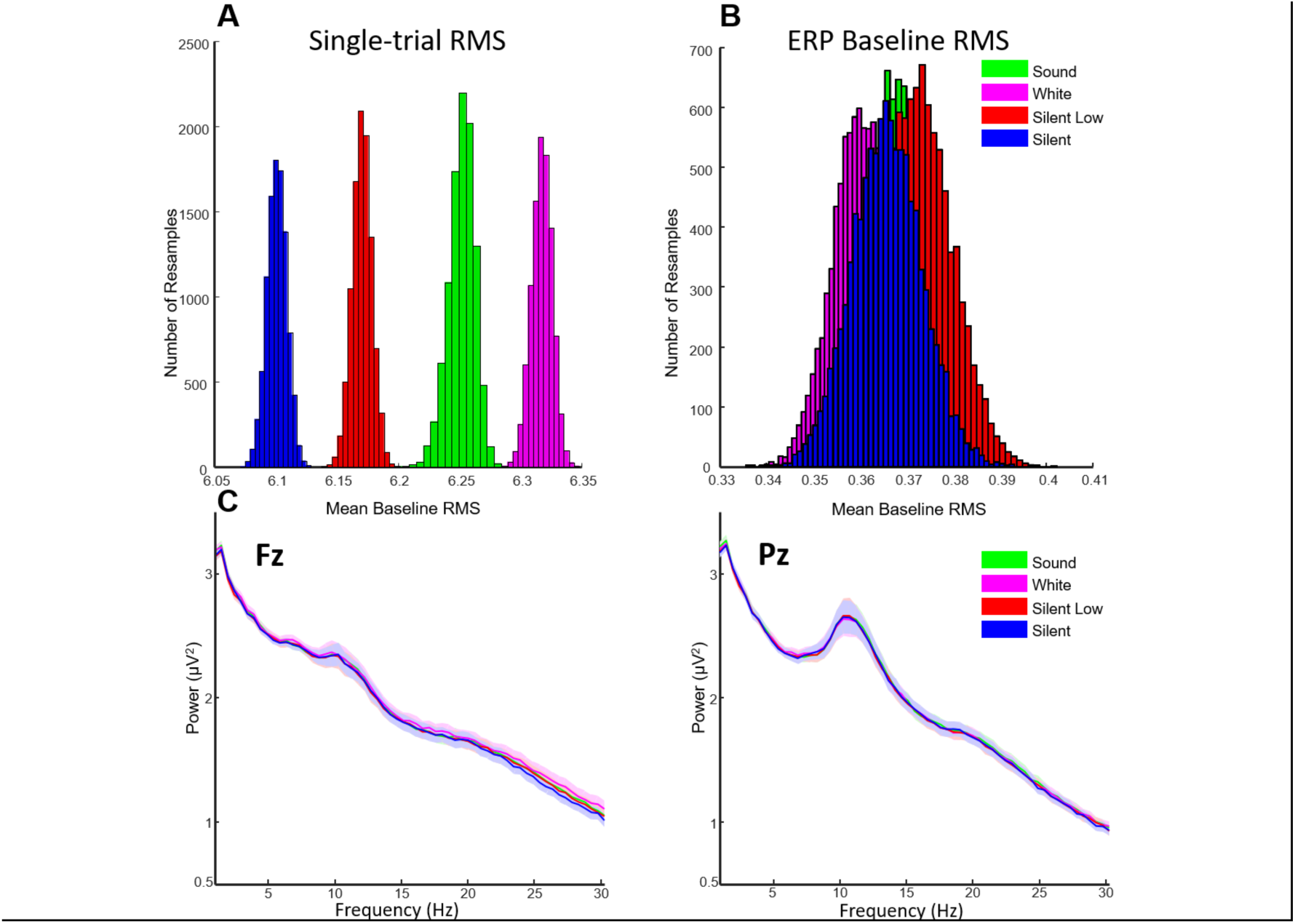
Difference waves and spectral analysis. A: Histogram of single-trial root mean square (RMS) grand average values collected during a 200 ms baseline period prior to tone onset, for 10,000 permutations of 360 randomly chosen standard trials for each subject. RMS values are averaged over all electrodes within each trial, then averaged over trials, and then averaged over subjects. B: Histogram of ERP baseline RMS values, calculated using 10,000 randomly selected permutations of 360 standard trials for each subject. RMS of the baseline period was computed for each permutation, and the data averaged over trials. C: Single-trial EEG spectra from electrodes Fz and Pz. This was calculated with zero-padded FFTs on 290 auditory standard trial epochs for each subject, averaged over trials, then subjects. Shaded regions indicate the standard error of the mean. Red asterisks denote significant differences, while green asterisks denote a marginal difference.

#### 3.5 Single-trial and ERP baseline noise

A grand average single-trial RMS was computed and recorded for each of these random selections and each condition. Figure 5A shows a histogram of the grand-averaged single-trial RMS values calculated for each permutation, for each condition. Evident from these plots is a small distinction between the single-trial noise between single-trial noise of the four conditions. A Wilcoxon signed rank test was used to compare each of the conditions (Table 3). The histogram in figure 5B shows the grand average RMS values calculated with these 10,000 permutations for each condition. A Wilcoxon signed rank test was then used to compare each of the conditions (Table 3).

**Table 2.** Mean and standard deviation of N1, P2 and P3 ERP components for both standard and target stimuli.

| ERP component mean and standard deviation |  |  |  |  |
| --- | --- | --- | --- | --- |
| Condition | Targets |  | Standards |  |
| N1 | Mean | Standard Deviation | Mean | Standard Deviation |
| Sound | -4.4634 | 2.1804 | -4.0088 | 2.0198 |
| White noise | -4.3358 | 2.2828 | -3.6449 | 2.3172 |
| Silent-low | -3.8537 | 2.1576 | -3.2266 | 2.1154 |
| Silent | -3.3709 | 2.6494 | -2.3262 | 2.2225 |
| P2 | Mean | Standard Deviation | Mean | Standard Deviation |
| Sound | -1.7616 | 2.9640 | -0.4898 | 2.0051 |
| White noise | -1.0464 | 3.0749 | 0.8429 | 2.3531 |
| Silent-low | -0.8887 | 3.0258 | 1.8717 | 1.9144 |
| Silent | 0.0235 | 3.3984 | 1.8582 | 2.2029 |
| P3 | Mean | Standard Deviation | Mean | Standard Deviation |
| Sound | 5.7235 | 4.7586 | -0.0156 | 1.3159 |
| White noise | 6.1974 | 4.4719 | -0.2612 | 1.0621 |
| Silent-low | 5.6705 | 4.4136 | -0.4813 | 1.1910 |
| Silent | 6.0389 | 4.7318 | 0.20211 | 0.8203 |

**Table 3.**
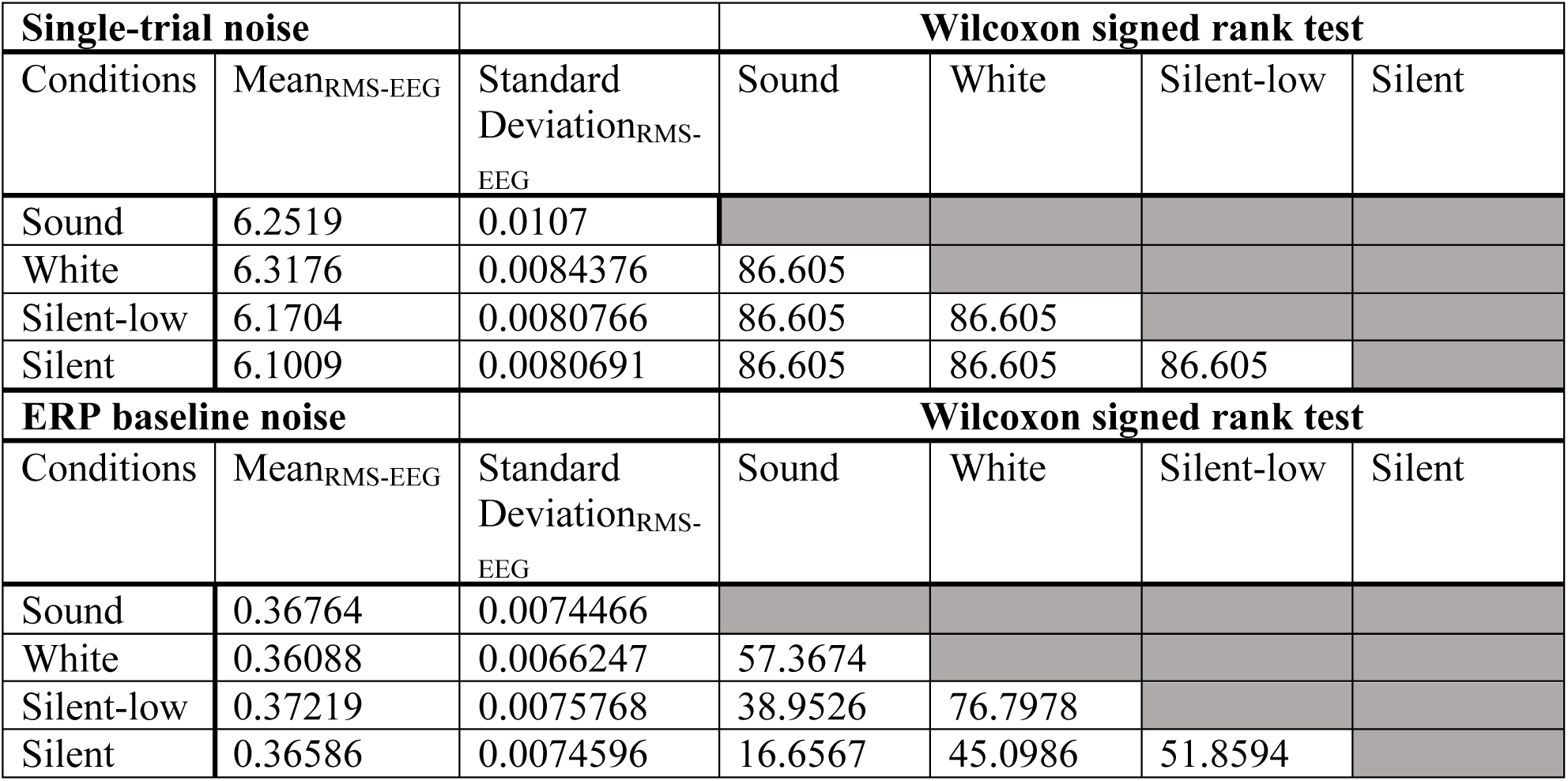
Mean and standard deviations of the single-trial and ERP baseline noise for each condition, as well as Wilcoxon signed rank test comparing all conditions.

| Single-trial noise |  |  | Wilcoxon signed rank test |  |  |  |
| --- | --- | --- | --- | --- | --- | --- |
| Conditions | Mean<sub>RMS-EEG</sub> | Standard Deviation<sub>RMS-EEG</sub> | Sound | White | Silent-low | Silent |
| Sound | 6.2519 | 0.0107 |  |  |  |  |
| White | 6.3176 | 0.0084376 | 86.605 |  |  |  |
| Silent-low | 6.1704 | 0.0080766 | 86.605 | 86.605 |  |  |
| Silent | 6.1009 | 0.0080691 | 86.605 | 86.605 | 86.605 |  |
| ERP baseline noise |  |  | Wilcoxon signed rank test |  |  |  |
| Conditions | Mean<sub>RMS-EEG</sub> | Standard Deviation<sub>RMS-EEG</sub> | Sound | White | Silent-low | Silent |
| Sound | 0.36764 | 0.0074466 |  |  |  |  |
| White | 0.36088 | 0.0066247 | 57.3674 |  |  |  |
| Silent-low | 0.37219 | 0.0075768 | 38.9526 | 76.7978 |  |  |
| Silent | 0.36586 | 0.0074596 | 16.6567 | 45.0986 | 51.8594 |  |

## 4. Discussion

In this study we compared ERPs during an oddball task in four conditions: with background traffic sounds (*outdoor sounds*), background white noise (*white noise*), silent background with quiet tones (*silent-low*), and silent background (*silent*). We found that we were able to replicate the main findings of a previous study (Scanlon et al., 2017b) in which an increased N1 and decreased P2 were observed while participants biked outside. In particular, when we played a recording of *outdoor sounds* in the background while participants performed a headphone auditory oddball task, the N1 was increased and the P2 was decreased compared to the *silent* condition. This indicates that these sounds had an effect on the sensory processing involved with the auditory task. The present paper was also able to look at reaction time and response rate, finding no behavioural differences between the conditions. Additionally, the *white noise* condition had a similar effect to a lesser extent, indicating that a simple, unchanging background sound can have a similar effect on sensory processes. Finally, the *silent-low* condition, in which the tones themselves were played at a quieter volume also appeared to have an effect, but only at the N1, indicating that a quiet task volume may also have some effect on these sensory processes. While we cannot say for certain that no other factors played a part in the N1-P2 effect found in the first study (Scanlon et al., 2017b), we can determine that background noises were at least sufficient to replicate this effect.

### 4.1 Alpha power

We investigated changes in alpha power for the differing sound conditions because previous studies have indicated that alpha power can influence N1 and P2 amplitudes (Brandt, Jansen & Carbonari, 1991; Jansen, & Brandt, 1991). However, in this study, no differences in alpha power were found between conditions, demonstrating that the effects within the N1 and P2 were independent of alpha power. A previous mobile EEG study from this lab did show a marginal difference in alpha power while participants cycled outside compared to sitting inside (Scanlon et al., 2017b), however this is presumably due to the increase in awareness to visual stimuli present while participants cycled outside. Alpha is known to reflect an individual’s state of awareness (Mathewson et al., 2011), as it has been demonstrated to increase with decreasing attentional focus, with the highest power when participants’ eyes were closed (Berger, 1929; Adrian & Matthews, 1934). When participants perform a task outside they are drawn to attend to their environment, while indoor experiments including the present study do not have this effect.

### 4.2 The N1 component

According to Näätänen & Picton (1987), auditory N1 can be functionally and topographically separated into up to six components. One of these components is known for an attentional function of sensory acceptance and rejection, which appears to amplify auditory perception of interesting, pleasant or important stimuli while attenuating the perception of uninteresting, unpleasant or unimportant stimuli (Näätänen & Picton, 1987; Roder et al., 1999). This N1 component has been referred to as an early, rapid selection of competing auditory ‘channels’ or auditory inputs, defined by differing pitch or location cues (Hansen & Hillyard, 1988). For example, the N1 tends to be increased when a sensory stimulus is located where one was already attending, and decreased when the same stimulus appears anywhere else, (Teder-Sälejärvi, et al. 1998). This is believed to allow enhanced processing of the stimuli of interest, and the effect is increased when the individuals performing the task are blind (Roder et al., 1999). The N1 component tends to have shorter latency and larger amplitude when the attended and unattended sounds are easily distinguishable by physical cues such as their pitch or location (Näätänen, 1982, 1992). This component has been shown to have increased amplitude as a function of increased attentional allocation to a specific input channel, as well as more accurate behavioural target detection for targets in that channel (Hink, Voorhis, Hillyard, & Smith, 1977). Functionally, the N1 amplitude and latency is believed to represent the amount of sensory information moving through an early mechanism of channel selection (Hillyard, Hink, Schwent, & Picton, 1973), further processing of information from the attended channel (Näätänen, 1982; Okita, 1981) as well as how well the eliciting stimulus and cue characteristics match within the attended input channel (Näätänen, 1992). In this study, it appears that the N1 increased when participants had to pay more attention and ‘tune in’ in order to perform the task properly in less than ideal conditions, and therefore was increased when *outdoor sounds* and *white noise* were played in the background, as well in the *silent-low* condition when the tones were simply quieter.

Another component of the N1 is also altered simply by stimulus intensity (Folsom & Owsley, 1987; Bruneau, Roux, Guerin, Garreau, Lelord, 1992; Näätänen & Picton, 1987) in that it is usually increased with increasing stimulus intensity, or volume. This contradicts the present study, as we found a significant increase in N1 amplitude when the stimulus had a decreased intensity (e.g. in the *silent low* condition). However this tendency to increase with increasing stimulus intensity is subject to inter-individual variability and N1 amplitude can, sometimes be decreased with increased stimulus intensity (Bruneau, et al., 1992), or even be leveled-off at high intensities (Buchsbaum, 1976). According to the ‘stimulus intensity’ interpretation, it is possible to interpret the N1 increases in the outside sound and white noise conditions as simply due to an increase in overall stimulus intensity during the task. However, few studies have used background noise or concurrent sounds as a manipulation for the N1, and those that have tend to show decreased N1 amplitude when concurrent sounds were present (Folsom & Owsley, 1987; Getzman, Golob & Wascher, 2016). It is also possible that the N1 amplitude increase in the *silent low* condition happened for an entirely different reason than the N1 amplitude increase in the *outdoor sounds* and *white noise* conditions, or even a combination of reasons that could also include arousal, context or inter-subject variability. With the current design, it is not possible to determine the exact reason for the changes in N1 amplitude between sound conditions. At this moment, we can only conclude that the N1 was altered by changes in sound stimuli, and that in our previous studies this could not have been entirely caused by outdoor activity or visual stimuli.

### 4.3 The P2 component

P2 amplitude has been shown to decrease to attended stimuli (Crowley & Colrain, 2004). The P2 has been found to reflect stimulus evaluation and classification as well as attentional allocation (Potts, 2004). For example, this component has been shown to decrease in response to speech sounds in a cocktail party task, while individuals had to pay attention to certain speech cues while ignoring simultaneous irrelevant speech sounds (Getzman, Golob & Wascher, 2016). This may be related to a mechanism of how effectively stimuli are able to be discriminated, as several studies have shown that the auditory P2 increases in amplitude after discrimination training (Atienza, Cantero, Dominguez-Marin, 2002; Hayes, Warrier, Nicol, Zecker, Kraus, 2003; Reinke, He, Wang, Alain, 2003; Trainor, Shahin, Roberts, 2003; Tong, Melara & Rao, 2009; Tremblay, Kraus, Carrell & McGee, 1997; Tremblay & Kraus, 2003) as well as with simple exposure to the stimulus (Sheehan, McArthur & Bishop, 2005). In our study, playing a conflicting sound in the background may have decreased the participant’s ability to discriminate the tones from background sounds in the *outdoor sounds* and *white noise* conditions, causing a decrease in the P2. This may also explain why there was no P2 effect in the quiet tones condition, as the lower volume did not decrease one’s ability to discriminate the tones. Further, salient and/or ‘eventful’ background stimuli may further amplify this effect as the *outdoor sounds* had a significantly smaller P2 amplitude than the *white noise* condition within standard tones.

As no behavioural differences were found, it appears that both the N1 and P2 represent two separate but related processes that allow individuals to perform the task sufficiently in a noisy environment. These hypotheses require future investigations both inside and outside the lab.

### 4.3 Single-trial and ERP baseline noise

Single trial and ERP baseline RMS noise were also affected by sound conditions, as shown in figure 5A and B and Table 3. Here we can see that the consistent background sound during the *white noise* condition was translated to a higher single-trial RMS data noise than any of the other conditions. We can also see, however, that this single-trial noise was averaged out over trials as the *white noise* condition has the lowest average ERP baseline noise. Additionally the plots show that the *outdoor sound* condition had the second-largest amount of both single-trial and ERP data noise, demonstrating that the inconsistency of this background noise may have decreased the ability for the data noise to be averaged out over trials. These small differences are an important consideration in terms of auditory experiments, as they show the ways in which sound can influence data noise. However in comparison to previous papers from this lab, the effect of sound conditions on RMS pre-stimulus data noise are at most approximately a quarter of the magnitude of those shown during outdoor cycling (Scanlon et al., 2017b) and half of those shown during stationary cycling (Scanlon et al., 2017a). Additionally, these effects did not appear to significantly carry over to any differences in spectral analysis (Figure 5C).

### 4.4 Limitations

There are a few aspects of the present study that limit our conclusions. The sample size of 14 subjects is relatively low. However, the study is based on similar previous studies with a sample size of 12 (Scanlon et al., 2017b) and 14 (Scanlon et al., 2017a), which both had considerably more data noise due to movement conditions and were able to show results within the ERP components selected for this study. Therefore, we believe that this study has the same ability to present accurate results. Indeed, our clear P3 effects in the current study show a high degree of power needed to measure ERP effects reliably. Given the high degree of within subject precision due to high number of trials and consistent trials, classic measures of power are less appropriate to estimate reliability of results (Luck, 2019). Another issue is that while the study was planned to have equally counterbalanced condition order, two participants were removed during analysis due to technical and task performance problems, causing the counterbalancing to be less than ideal. This misbalanced counterbalancing is particularly non-ideal in light of a relatively low sample-size. However, if condition order effects were present one would expect to see behavioural effects within the reaction time data due to practice effects, while in the present study there were no behavioural differences between conditions.

### 4.5 Future Directions

With this new knowledge about the functions of the N1 and P2 components, we intend to investigate further into how these components relate to the way humans perform auditory and visual tasks in ecologically valid environments. In particular, an interesting observation from this paper is that the P2 appears to be somewhat related proportionally to the volume of background noise, being decreased in amplitude with increasing background noise. One future direction might be to play comparably similar background noises at different volumes during another auditory task, to see if the P2 can be directly correlated with volume of the background noise. Additionally, our lab intends to perform studies on the visual ERP when the task is performed with a visually noisy background, as well as cross modal studies in which an auditory task is used in a visually noisy environment, and vice versa. As well, we plan to perform further outdoor studies in which auditory and visual tasks are performed in environments with differing auditory and visual noise. The main goal is to examine exactly how the human brain is able to compensate and perform tasks effectively in the face of distracting stimuli.

### 4.6 Conclusion

In this study, we replicated the effects found in a previous outdoor ERP study of modulated early sensory ERP components by only replaying the background sounds from the previous environment. *Outdoor traffic sounds* and *white noise* both modulated the N1 and P2 components of the ERPs evoked during an auditory oddball task. We found evidence that the N1 and P2, while functionally related, appear to perform two distinct mechanisms during stimulus discrimination in ecologically valid conditions. It appears that the N1 was altered by stimulus changes between the conditions, possibly increasing with attentiveness to a particular channel of auditory input, while the P2 decreased with attention and as the auditory channel became more difficult to discriminate. This effect may be part of a mechanism that allows one to selectively attend to auditory tasks in noisy sound environments, as humans often experience in day to day life.

## References

Adrian, E. D., and Matthews, B. H. C. (1934). The Berger rhythm: potential changes from the occipital lobes in man. Brain 4, 355–385.

Atienza, M., Cantero, J. L., & Dominguez-Marin, E. (2002). The time course of neural changes underlying auditory perceptual learning. Learning & Memory, 9(3), 138–150.

Ballas, J. A., & Howard, J. H. (1987). Interpreting the language of environmental sounds. Environment and behavior, 19(1), 91–114.

Beagley, H., & Knight, J. (1967). Changes in auditory evoked response with intensity. The Journal of Laryngology & Otology, 81(08), 861–873. doi:https://doi.org/10.1017/S0022215100067815

Berger, H. (1929). Über das elektrenkephalogramm des menschen. Archiv für psychiatrie und nervenkrankheiten, 87(1), 527–570.

Brandt, M. E., Jansen, B. H., & Carbonari, J. P. (1991). Pre-stimulus spectral EEG patters and the visual evoked response. Electroencephalography and Clinical Neurophysiology/Evoked Potentials Section, 80(1), 16–20. doi:https://doi.org/10.1016/0168-5597(91)90037-X

Bruneau, N., Roux, S., Guérin, P., Garreau, B., & Lelord, G. (1993). Auditory stimulus intensity responses and frontal midline theta rhythm. Electroencephalography and Clinical Neurophysiology, 86(3), 213–216.

Buchsbaum, M. (1976). Self-regulation of stimulus intensity: Augmenting/reducing and the average evoked response. In Consciousness and self-regulation (pp. 101–135). Springer, Boston, MA.

Crowley, K. E., & Colrain, I. M. (2004). A review of the evidence for P2 being an independent component process: age, sleep and modality. Clinical neurophysiology, 115(4), 732–744. doi:https://doi.org/10.1016/j.clinph.2003.11.021

Debener, S., Minow, F., Emkes, R., Gandras, K., & de Vos, M. (2012). How about taking a low-cost, small, and wireless EEG for a walk? Psychophysiology, 49(11), 1449–1453. doi:10.1111/j.1469-8986.2012.01471.x

De Vos, M., Gandras, K., & Debener, S. (2014). Towards a truly mobile auditory brain– computer interface: exploring the P300 to take away. International journal of psychophysiology, 91(1), 46–53. doi:https://doi.org/10.1016/j.ijpsycho.2013.08.010

Dunn, B. R., Dunn, D. A., Languis, M., & Andrews, D. (1998). The relation of ERP components to complex memory processing. Brain and cognition, 36(3), 355–376. doi:https://doi.org/10.1006/brcg.1998.0998

Federmeier, K. D., & Kutas, M. (2002). Picture the difference: Electrophysiological investigations of picture processing in the two cerebral hemispheres. Neuropsychologia, 40(7), 730–747. doi:https://doi.org/10.1016/S0028-3932(01)00193-2

Freunberger, R., Klimesch, W., Doppelmayr, M., & Höller, Y. (2007). Visual P2 component is related to theta phase-locking. Neuroscience letters, 426(3), 181–186. doi:https://doi.org/10.1016/j.neulet.2007.08.062

Folsom, R. C., & Owsley, R. M. (1987). N1 action potentials in humans: influence of simultaneous contralateral stimulation. Acta oto-laryngologica, 103(3-4), 262–265.

Getzmann, S., Golob, E. J., & Wascher, E. (2016). Focused and divided attention in a simulated cocktail-party situation: ERP evidence from younger and older adults. Neurobiology of aging, 41, 138–149. doi:https://doi.org/10.1016/j.neurobiolaging.2016.02.018

Griffiths, T. D., Warren, J. D. (2004). What is an auditory object? Nature Reviews Neuroscience, 5, 887–892. doi:https://doi.org/10.1038/nrn1538

Hackley, S. A., Woldorff, M., & Hillyard, S. A. (1990). CrossnModal Selective Attention Effects on Retinal, Myogenic, Brainstem, and Cerebral Evoked Potentials. Psychophysiology, 27(2), 195–208. doi:https://doi.org/10.1111/j.1469- 8986.1990.tb00370.x

Hansen, J. C., & Hillyard, S. A. (1988). Temporal dynamics of human auditory selective attention. Psychophysiology, 25(3), 316–329.

Hansenne, M. (2000). Le potentiel évoqué cognitif P300 (I): aspects théorique et psychobiologique. Neurophysiologie Clinique/Clinical Neurophysiology, 30(4), 191–210. doi:https://doi.org/10.1016/S0987-7053(00)00223-9

Hayes, E. A., Warrier, C. M., Nicol, T. G., Zecker, S. G., & Kraus, N. (2003). Neural plasticity following auditory training in children with learning problems. Clinical neurophysiology, 114(4), 673–684. doi:https://doi.org/10.1016/S1388-2457(02)00414-5

Hillyard, S. A, Hink, R. F, Schwent, V. L., & Picton, T. W. (1973). Electrical signs of selective attention in the human brain. Science, 182(4108), 177–180. doi:https://doi.org/10.1126/science.182.4108.177

Hink, R. F, Voorhis, S. T. Van Hillyard, S. A., & Smith, T. (1977). The division of attention and the human auditory evoked potentials. Neuropsychologia, 15(4-5), 597–605. doi:https://doi.org/10.1016/0028-3932(77)90065-3

Jansen, B. H., & Brandt, M. E. (1991). The effect of the phase of prestimulus alpha activity on the averaged visual evoked response. Electroencephalography and Clinical Neurophysiology/Evoked Potentials Section, 80(4), 241–250. doi:https://doi.org/10.1016/0168-5597(91)90107-9

Krigolson, O. E., Williams, C. C., Norton, A., Hassall, C. D., & Colino, F. L. (2017). Choosing MUSE: Validation of a low-cost, portable EEG system for ERP research. Frontiers in neuroscience, 11, 109.

Kirmse, U., Jacobsen, T., & Schröger, E. (2009). Familiarity affects environmental sound processing outside the focus of attention: An event-related potential study. Clinical neurophysiology, 120(5), 887–896. doi:https://doi.org/10.1016/j.clinph.2009.02.159

Kuziek, J. W., Shienh, A., Mathewson, K. E. (2017). Transitioning EEG experiments away from the laboratory using a Raspberry Pi 2. Journal of Neuroscience Methods, 277, 75–82. doi:https://doi.org/10.1016/j.jneumeth.2016.11.013

Lefebvre, C. D., Marchand, Y., Eskes, G. A., & Connolly, J. F. (2005). Assessment of working memory abilities using an event-related brain potential (ERP)-compatible digit span backward task. Clinical Neurophysiology, 116(7), 1665–1680. doi:https://doi.org/10.1016/j.clinph.2005.03.015

Loftus, G. R., & Masson, M. E. (1994). Using confidence intervals in within-subject designs. Psychonomic bulletin & review, 1(4), 476–490. http://dx.doi.org/10.3758/BF03210951

Luck, S. J., & Hillyard, S. A. (1994). Electrophysiological correlates of feature analysis during visual search. Psychophysiology, 31(3), 291–308

Luck, S. J. (2014). An introduction to the event-related potential technique. Cambridge, MA: MIT Press. doi:10.1086/506120

Luck, S. J. (2019, February 20). Why experimentalists should ignore reliability and focus on precision. Retrieved March 29, 2019, from https://lucklab.ucdavis.edu/blog/2019/2/19/reliability-and-precision

Mathewson, K. E., Lleras, A., Beck, D. M., Fabiani, M., Ro, T., & Gratton, G. (2011). Pulsed out of awareness: EEG alpha oscillations represent a pulsed-inhibition of ongoing cortical processing. Frontiers in psychology, 2, 99.

Mathôt, S., Schreij, D., & Theeuwes, J. (2012). OpenSesame: An open-source, graphical experiment builder for the social sciences. Behavior research methods, 44(2), 314–324.

Näätänen, R., & Picton, T. (1987). The N1 wave of the human electric and magnetic response to sound: a review and an analysis of the component structure. Psychophysiology, 24(4), 375–425. doi:https://doi.org/10.1111/j.1469-8986.1987.tb00311.x

Näätänen, R. (1982). Processing negativity: An evoked-potential reflection of selective attention. Psychological Bulletin, 92, 605–640. doi:http://dx.doi.org/10.1037/0033-2909.92.3.605

Näätänen, R. (1992). Attention and brain function. Hillsdale, NJ: Erlbaum.

Okita, T. (1981). Slow negative shifts of the human event-related potential associated with selective information processing. Biological Psychology, 12(1), 63–75. doi:https://doi.org/10.1016/0301-0511(81)90020-X

Paiva, T. O., Almeida, P. R., Ferreira-Santos, F., Vieira, J. B., Silveira, C., Chaves, P. L., et al. (2016). Similar sound intensity dependence of the N1 and P2 components of the auditory ERP: Averaged and single trial evidence. Clinical Neurophysiology, 127(1), 499–508. doi:https://doi.org/10.1016/j.clinph.2015.06.016

Picton, T., Goodman, W., & Bryce, D. (1970). Amplitude of evoked responses to tones of high intensity. Acta oto-laryngologica, 70(2), 77–82. doi:https://doi.org/10.3109/00016487009181862

Potts, G. F. (2004). An ERP index of task relevance evaluation of visual stimuli. Brain and cognition, 56(1), 5–13. doi:https://doi.org/10.1016/j.bandc.2004.03.006

Rapin, I., Schimmel, H., Tourk, L. M., Krasnegor, N. A., & Pollak, C. (1966). Evoked responses to clicks and tones of varying intensity in waking adults. Electroencephalography and clinical neurophysiology, 21(4), 335–344. doi:https://doi.org/10.1016/0013-4694(66)90039-3

Reinke, K. S., He, Y., Wang, C., & Alain, C. (2003). Perceptual learning modulates sensory evoked response during vowel segregation. Cognitive Brain Research, 17(3), 781–791. doi:https://doi.org/10.1016/S0926-6410(03)00202-7

Röder, B., Teder-Sälejärvi, W., Sterr, A., Rösler, F., Hillyard, S. A., & Neville, H. J. (1999). Improved auditory spatial tuning in blind humans. Nature, 400(6740), 162.

Roye, A., Jacobsen, T., & Schröger, E. (2013). Discrimination of personally significant from nonsignificant sounds: A training study. Cognitive, Affective, & Behavioral Neuroscience, 13(4), 930–943. doi:https://doi.org/10.3758/s13415-013-0173-7

Scanlon, J. E. M., Sieben, A. J., Holyk, K. R., & Mathewson, K. E. (2017a). Your brain on bikes: P3, MMN/N2b, and baseline noise while pedaling a stationary bike. Psychophysiology, 54(6), 927–937. doi:https://doi.org/10.1111/psyp.12850

Scanlon, J. E. M., Townsend, K. A., Cormier, D. L., Kuziek, J. W., & Mathewson, K. E. (2017b). Taking off the training wheels: Measuring auditory P3 during outdoor cycling using an active wet EEG system. Brain research. doi:https://doi.org/10.1016/j.brainres.2017.12.010

Shahin, A., Bosnyak, D. J., Trainor, L. J., & Roberts, L. E. (2003). Enhancement of neuroplastic P2 and N1c auditory evoked potentials in musicians. The Journal of Neuroscience, 23(13), 5545–5552. doi:https://doi.org/10.1523/JNEUROSCI.23-13-05545.2003

Shahin, A. J., Roberts, L. E., Miller, L. M., McDonald, K. L., & Alain, C. (2007). Sensitivity of EEG and MEG to the N1 and P2 auditory evoked responses modulated by spectral complexity of sounds. Brain topography, 20(2), 55–61. doi:https://doi.org/10.1007/s10548-007-0031-4

Shahin, A., Roberts, L. E., Pantev, C., Trainor, L. J., & Ross, B. (2005). Modulation of P2 auditory-evoked responses by the spectral complexity of musical sounds. Neuroreport, 16(16), 1781–1785. doi:https://doi.org/10.1097/01.wnr.0000185017.29316.63

Sheehan, K. A., McArthur, G. M., & Bishop, D. V. (2005). Is discrimination training necessary to cause changes in the P2 auditory event-related brain potential to speech sounds? Cognitive Brain Research, 25(2), 547–553. doi:https://doi.org/10.1016/j.cogbrainres.2005.08.007

Teder-Sälejärvi, W. A., & Hillyard, S. A. (1998). The gradient of spatial auditory attention in free field: an event-related potential study. Perception & Psychophysics, 60(7), 1228-1242.

Thaerig, S., Behne, N., Schadow, J., Lenz, D., Scheich, H., Brechmann, A., et al. (2008). Sound level dependence of auditory evoked potentials: simultaneous EEG recording and low-noise fMRI. International Journal of Psychophysiology, 67(3), 235–241. doi:https://doi.org/10.1016/j.ijpsycho.2007.06.007

Trainor, L. J., Shahin, A., & Roberts, L. E. (2003). Effects of musical training on the auditory cortex in children. Annals of the New York Academy of Sciences, 999(1), 506–513. doi:https://doi.org/10.1196/annals.1284.061

Tremblay, K., Kraus, N., Carrell, T. D., & McGee, T. (1997). Central auditory system plasticity: generalization to novel stimuli following listening training. The Journal of the Acoustical Society of America, 102(6), 3762–3773. doi:https://doi.org/10.1121/1.420139

Tremblay, K. L., & Kraus, N. (2002). Auditory training induces asymmetrical changes in cortical neural activity. Journal of Speech, Language, and Hearing Research, 45(3), 564–572. doi:doi:10.1044/1092-4388(2002/045)

Tremblay, K., Kraus, N., McGee, T., Ponton, C., & Otis, B. (2001). Central auditory plasticity: changes in the N1-P2 complex after speech-sound training. Ear and Hearing, 22(2), 79– 90.

Tong, Y., Melara, R. D., & Rao, A. (2009). P2 enhancement from auditory discrimination training is associated with improved reaction times. Brain research, 1297, 80–88. doi:https://doi.org/10.1016/j.brainres.2009.07.089

Verleger, R. (1988). Event-related potentials and cognition: A critique of the context updating hypothesis and an alternative interpretation of P3. Behavioral and Brain Sciences, 11(03), 343–356.

Wolpaw, J. R., & Penry, J. K. (1975). A temporal component of the auditory evoked response. Electroencephalography and clinical neurophysiology, 39(6), 609–620. doi:https://doi.org/10.1016/0013-4694(75)90073-5

Wood, C. C., & Wolpaw, J. R. (1982). Scalp distribution of human auditory evoked potentials. II. Evidence for overlapping sources and involvement of auditory cortex. Electroencephalography and Clinical Neurophysiology, 54(1), 25–38. doi:https://doi.org/10.1016/0013-4694(82)90228-0

Zink, R., Hunyadi, B., Van Huffel, S., & De Vos, M. (2016). Mobile EEG on the bike: Disentangling attentional and physical contributions to auditory attention tasks. Journal of Neural Engineering, 13(4), 046017. doi:doi:10.1088/1741-2560/13/4/046017

